# Attenuated amiloride-sensitive current and augmented calcium-activated chloride current in marsh rice rat (*Oryzomys palustris*) airways

**DOI:** 10.1101/624320

**Authors:** Shin-Ping Kuan, Yan-Shin J. Liao, Katelyn M. Davis, Jonathan G. Messer, Jasenka Zubcevic, J. Ignacio Aguirre, Leah R. Reznikov

**Author notes:** Contact Information: Leah Reznikov PhD, University of Florida, Department of Physiological Sciences, 1333 Center Drive, PO Box 100144, Gainesville FL 32610.

## Abstract

Prolonged heat and sea salt aerosols pose a challenge for the mammalian airway, placing the protective airway surface liquid (ASL) at risk for desiccation. Thus, mammals inhabiting salt marshes might have acquired adaptations for ASL regulation. We studied the airways of the rice rat, a rodent that inhabits salt marshes. We discovered negligible Na^+^ transport through the epithelial sodium channel (ENaC). In contrast, carbachol induced a large Cl^−^ secretory current that was blocked by the calcium-activated chloride channel (CaCC) inhibitor CaCCinh-A01. Decreased mRNA expression of α, β, and γ ENaC, and increased mRNA expression of the CaCC transmembrane member 16A distinguished the rice rat airway. Rice rat airway cultures also secreted fluid in response to carbachol and displayed an exaggerated expansion of the ASL volume when challenged with 3.5% NaCl. These data suggest that the rice rat airway might possess unique ion transport adaptations to facilitate survival in the salt marsh environment.

## Introduction

The airway epithelium is a critical barrier to the external environment and is protected by a thin layer of fluid known as the airway surface liquid (ASL). ASL facilitates the entrapment of particles and bacteria, prevents airway desiccation, humidifies inspired air, and supports mucociliary transport (Widdicombe, 1997). ASL volume and composition are regulated through active and passive epithelial cell ion transport that is in part dependent upon the apical epithelial sodium channel (ENaC) (Chambers et al., 2007), the cystic fibrosis transmembrane conductance regulator (CFTR) and calcium (Ca^2+^)-activated Cl^−^ channels (CaCC) (Frizzell and Hanrahan, 2012).

The importance of these proteins in regulating ASL composition and volume has been repeatedly shown. For example, in mice genetically modified to hyperabsorb Na^+^ through overexpression of the β ENaC subunit, ASL is depleted (Mall et al., 2004), whereas mice lacking the α ENaC subunit die shortly after birth due to an inability to clear fluid from their lungs (Hummler et al., 1996). Mutations in CFTR cause the life-shortening autosomal recessive disorder, cystic fibrosis (CF) (Riordan et al., 1989). The loss of CFTR-mediated Cl^−^ and bicarbonate secretion in CF sets up an airway environment prone to infection, inflammation, mucus obstruction, ASL depletion, and impaired fluid secretion (Boucher, 2007; Pezzulo et al., 2012). CaCC-mediated Cl^−^ transport also regulates ASL volume acutely (Tarran et al., 2002), and in mice lacking the CaCC, TMEM16A, there is defective fluid secretion (Rock et al., 2009).

Salt marshes are coastal wetlands characterized by high salt concentrations, intense heat, and periods of flooding (Silliman, 2014). The increased salinity and extreme temperatures pose a challenge for the mammalian airway. First, the dissipation of heat through panting (e.g., mouth breathing) can cause evaporative loss of water from the ASL lining the trachea and bronchi (Widdicombe, 1997). Second, inhalation of hypertonic solutions, such as those encountered in atmospheric sea-salt aerosols (Gabriel, 2002), can modify ASL composition. Both of these events can acutely increase ASL tonicity. The airway responds to hypertonic solutions via two major mechanisms. First, hypertonic solutions osmotically draw water onto the airway surface (Chambers et al., 2007; Harvey et al., 2011). Second hypertonic solutions activate the vagus nerve, causing the release of acetylcholine from nerve terminals innervating the airway (Wine, 2007). Acetylcholine then binds to muscarinic receptors located on epithelial cells, leading to increased intracellular Ca^2+^ and fluid secretion that is dependent upon activation of CaCC (Caputo et al., 2008; Catalan et al., 2015; Frizzell and Hanrahan, 2012).

The rice rat (*Oryzomys palustris*) is a medium-sized nocturnal rodent (40-80g) from the family *Cricetidae.* It can be found inhabiting the salt marshes of North and South America (Hamilton, 1946), where it is well-adapted for swimming (Wolfe and Esher, 1981). In captivity, the rice rat adapts well to standard laboratory conditions and requires minimal modifications (Aguirre et al., 2015).

In the current study, we hypothesized that the rice rat airway might have developed unique adaptations to ensure survival in the harsh salt marsh environment. Specifically, we hypothesized that exposure to high temperatures and sea salt aerosols might have resulted in an evolutionary pressure that decreased Na^+^ absorption through ENaC and increased dependence upon CaCC for Cl^−^ transport. To test this hypothesis, we examined net ion transport with Ussing chambers (Chen et al., 2010; Ostedgaard et al., 2011; Stoltz et al., 2013) and assessed fluid secretion in response to cholinergic stimulation and hypertonic solutions using reflective light microscopy (Harvey et al., 2011). Expression levels of key ion channels were also quantified. As controls, we utilized tracheal cultures of the mouse and the pig, animals that do not normally inhabit salt marshes and whose airway electrophysiological properties are well-characterized (Chen et al., 2010; Ostedgaard et al., 2011; Rogers et al., 2008; Stoltz et al., 2013).

## Results

### Cultured rice rat airways show exaggerated CaCC-mediated short circuit current, but negligible basal amiloride-sensitive short circuit current

We cultured primary tracheal airway epithelia from the rice rat, mouse and pig at an air-liquid interface. We first confirmed the presence of key features of airway epithelia, including cilia (Jain et al., 2010) (Figure 1A-C) and expression of the tight junction protein zonula occludens protein-1 (ZO-1) (Vermeer et al., 2009) (Figure 1D-1F). No staining was observed for either ZO-1 or acetylated alpha tubulin in negative controls (Figure 1G-L), suggesting specific detection.

**Figure 1.**
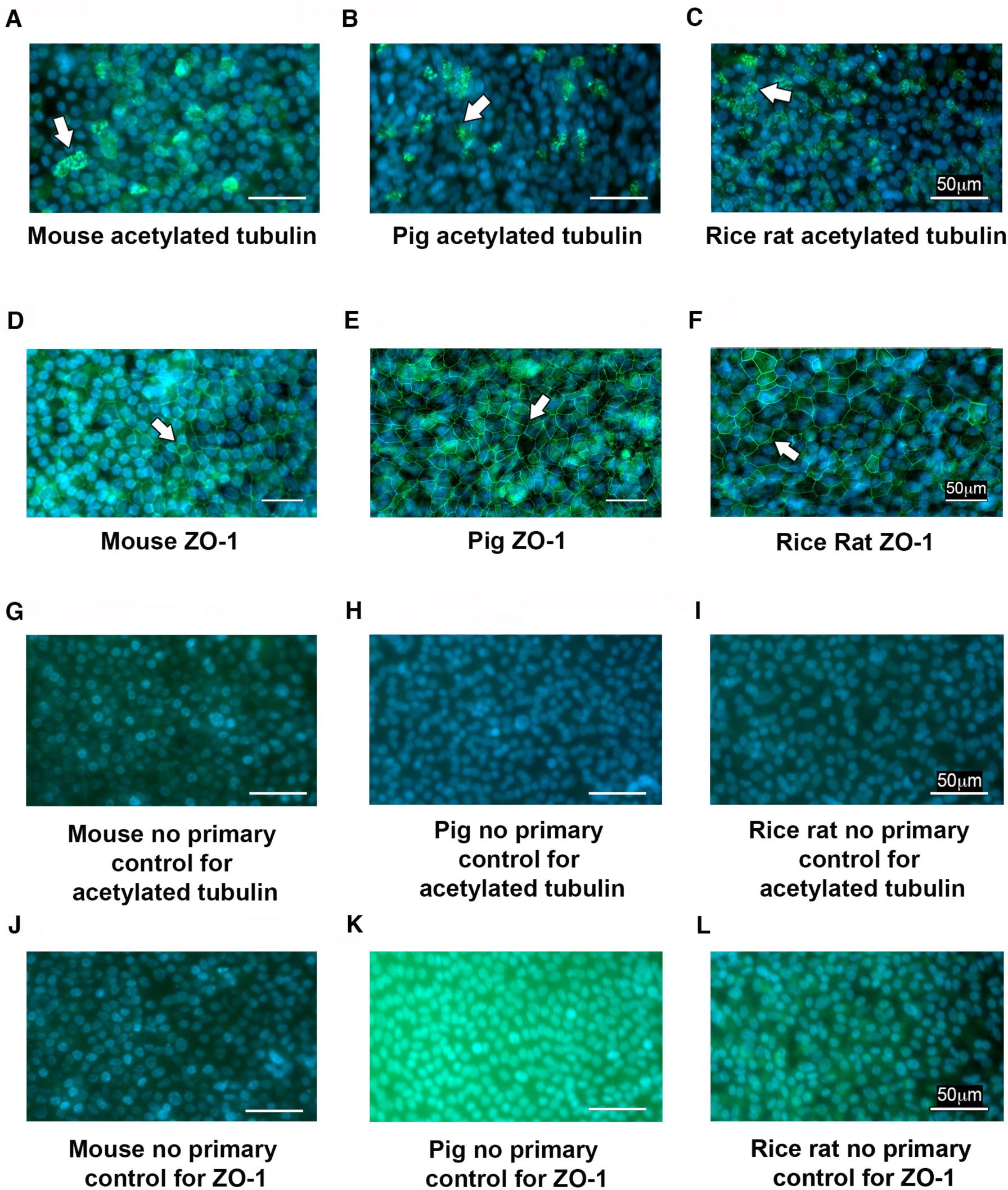
Tight junctions and cilia in rice rat cultures. En face images of tracheal cultures of mouse (A), piglet (B) and rice rat (C) showing expression of the cilia protein acetylated alpha tubulin. En face images of tracheal cultures of mouse (D), piglet (E) and rice rat (F) showing expression of the tight junction protein ZO-1. En face images of tracheal cultures of mouse (G), piglet (H) and rice rat (I) incubated with only the goat anti-mouse 488 secondary antibody (secondary used for the detection of acetylated tubulin). En face images of tracheal cultures of mouse (J), piglet (K) and rice rat (L) incubated with only goat anti-rabbit 488 secondary antibody; this secondary was used for the detection of ZO-1. Abbreviations: zonula occludens protein 1 (ZO-1). Hoechst was used to detect nuclei. For all panels, scale bars = 50 μm Antibody detection of ZO-1 and acetylated alpha tubulin was performed on two separate occasions for each species. White arrows show example of specific staining.

We next examined net ion transport. We found that the rice rat tracheal cultures displayed minimal basal short circuit current (Figure 2A, 2I). This basal value was significantly lower than the basal short circuit current measured in either the piglet or the mouse (Figure 2A, 2I-K). We also found minimal amiloride-sensitive Isc in the rice rat tracheal cultures compared to the mouse or pig (Figure 2B, 2I-K). These data suggested that the rice rat had diminished Na^+^ transport through ENaC.

**Figure 2.**
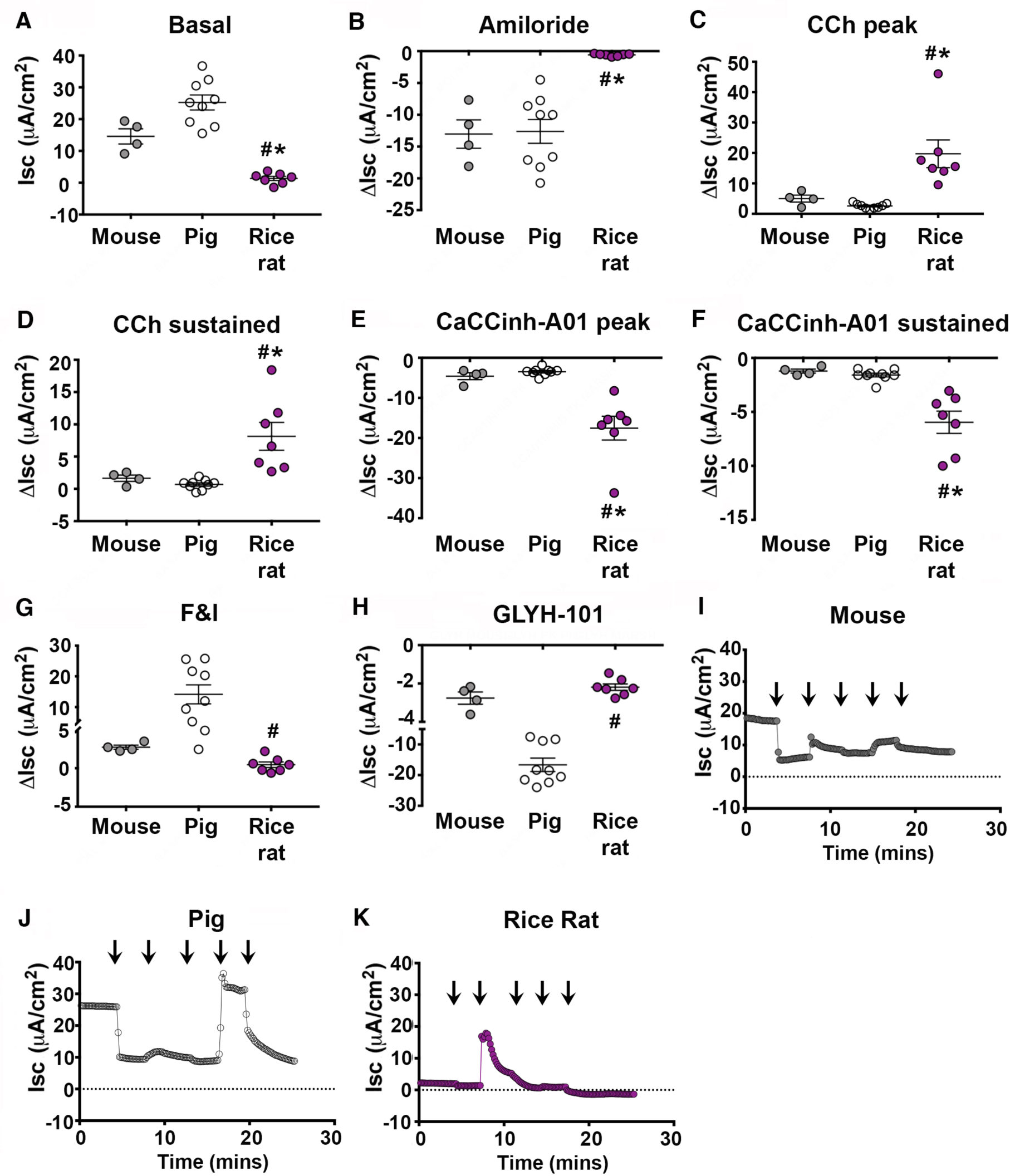
Primary tracheal cultures of rice rats have limited basal and amiloride-sensitive current, but large carbachol-mediated Cl^−^ secretion. (A) Basal short circuit current (Isc) measurements in primary cultures of rice rat and piglet. Averaged delta (Δ) Isc measurements shown for: 100μM apical amiloride (AMIL) (B); 100μM basolateral carbachol (CCh) (peak (C) and sustained (D) responses); 30μM apical CaCCinh-A01for inhibition of peak (E) and sustained (F) CCh-mediated current; apical 10μM forskolin and 100μM IBMX (F&I) (G), 100μM of apical GLYH-101 (H). Representative short circuit current (Isc) trace in mouse (I), piglet (J) and rice rat (K) airway cultures. Arrows indicate addition of drugs (as described in materials and methods). Abbreviations: carbachol (CCh); forskolin and IBMX (F&I). For all panels, n = 4 cultures from mice (2 female cultures and 2 male cultures representing 6 female and 6 male subjects); n = 7 cultures from the rice rat (4 female cultures, and 3 male cultures representing 8 female and 6 male subjects); n = 9 porcine cultures (5 female and 4 male representing 5 individual female and 4 individual male subjects. * = p < 0.05 compared to mouse; # = p < 0.05 compared to pig. All data are shown as mean ± SEM.

We next stimulated epithelium with carbachol basolaterally to activate calcium-activated chloride channels (CaCC) (Drumm et al., 1991). Carbachol induced a large transient outward current (Figure 2C) in rice rat cultures. Both the peak and sustained responses to carbachol were significantly greater in the rice rat compared to the mouse or pig (Figure 2C-D 2I-K). Application of CaCCinh-A01-inhibited carbachol-induced Cl^−^ secretion to a greater extent in the rice rat (Figure 2E-F, 2I-K). Thus, rice rat airways displayed augmented CaCC-mediated Cl^−^ secretion compared to pigs and mice.

We applied forskolin and IBMX (F&I) to activate CFTR. We found significantly less F&I-mediated Cl^−^ secretion in the rice rat compared to the pig, but not compared to the mouse (Figure 2G, 2I-K). Application of the CFTR inhibitor GlyH-101 caused a greater inhibition of F&I-mediated current in the pig compared to the rice rat (Figure 2H, 2J-K). The amount of GlyH-101-inhibited current in the rice rat and mouse was not different (Figure 2H, 2I, 2K). These data suggested that the rice rat had similar CFTR activity as the mouse, but comparatively less than the pig.

### Rice rat airway cultures have decreased expression of ENaC and increased expression of TMEM16a

Our ion transport studies suggested alterations in the transport mediated by ENaC and CaCC. Given that the rice rat genome is unknown, the tools available for quantifying ion channel expression at either the transcript or protein level are limited. To circumvent this, we cloned small segments of mRNA for rice rat CFTR, TMEM16A, α ENaC, β ENaC, γ ENaC, CHRM3 and RPL13a. Based upon those sequences, we then designed “universal” primers to allow for direct comparisons between mouse and rice rat using the same primer set. RPL13a was used as a housekeeping control (Shah et al., 2016). We preferentially focused on the mouse and rice rat because they are more closely related phylogenetically.

We found decreased mRNA expression of α, β and γ ENaC in the rice rat compared to the mouse (Figure 3A-C). To examine whether decreased mRNA expression might be due to a steroid hormone requirement, we treated rice rat cultures with aldosterone and dexamethasone using concentrations that augment ENaC expression in primary rat airway cells (Champigny et al., 1994). Twenty-four hours later, we re-examined α, β, and γ ENaC expression but found no effect (Figure 3D-F). Thus, these data suggested that rice rat did not share the same steroid requirement for ENaC expression as the rat.

**Figure 3.**
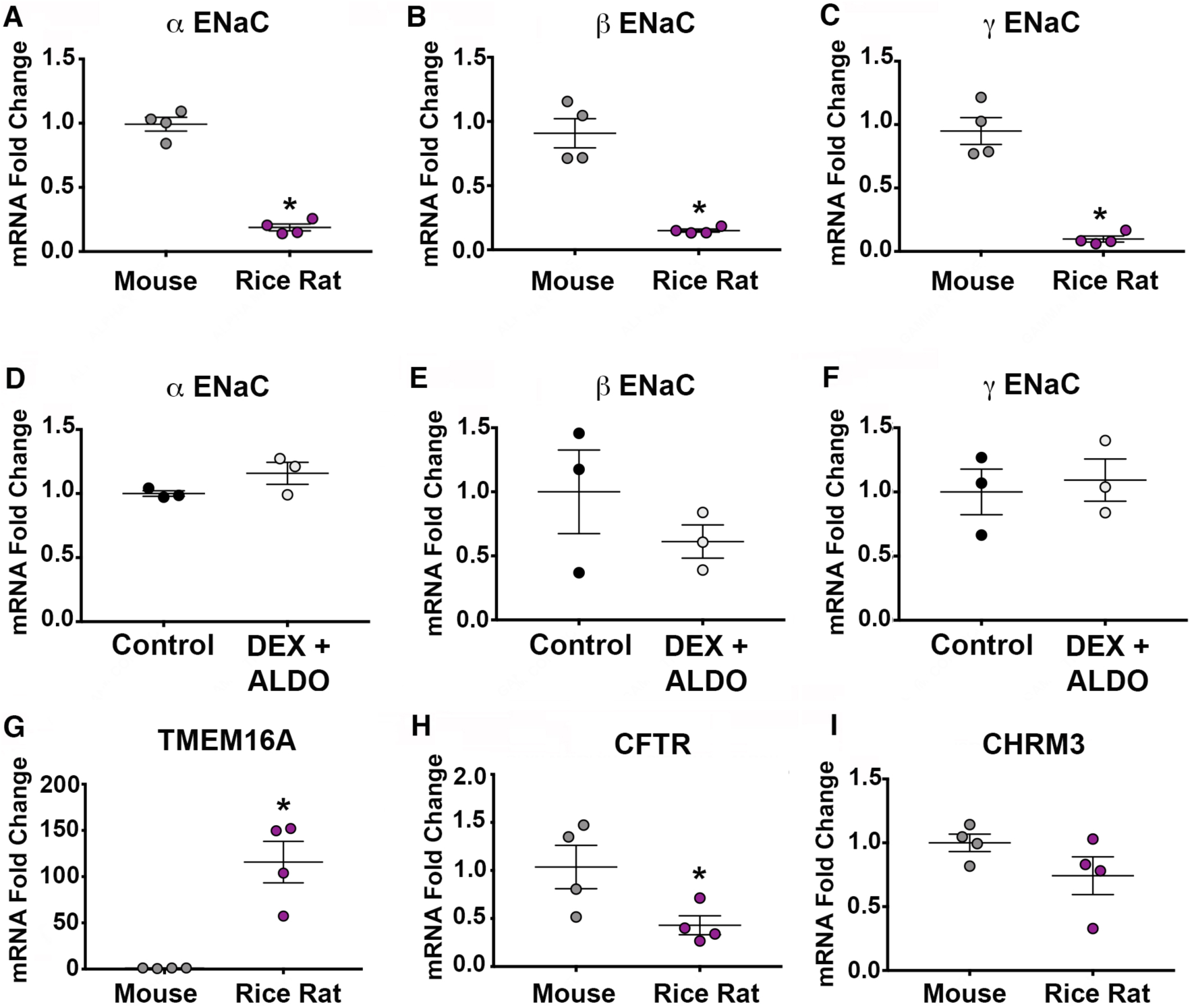
Rice rat tracheal cultures have decreased ENaC expression and increased TMEM16A expression compared to mice. mRNA expression of α ENaC (A), β ENaC (B), and γ ENaC (C) in mouse and rice rat airway cultures. mRNA expression of α ENaC (D), β ENaC (E), and γ EnaC (F) in rice rat airway cultures treated with dexamethasone (DEX) and aldosterone (ALDO). mRNA expression of CFTR (G), TMEM16A (H), and CHRM3 (I). Abbreviations: epithelial sodium channel (ENaC); cystic fibrosis transmembrane conductance regulator (CFTR); Transmembrane member 16A (TMEM16); muscarinic acetylcholine receptor 3 (CHRM3); dexamethasone (DEX); aldosterone (ALDO). For all panels A-C and G-I, n = 4 cultures from mice (2 female cultures and 2 male cultures representing 6 female and 6 male subjects); n = 4 cultures from the rice rat (2 female cultures, and 2 male cultures representing 6 female and 6 male subjects). For panels D-F, n = 3 cultures from the rice rat (2 female cultures, and 1 male culture representing 6 female and 3 male subjects). * = p < 0.05 compared to mouse. All data are shown as mean ± SEM.

We also examined transcript abundance of CFTR and TMEM16A in rice rat and mouse tracheal cultures. We found a robust increase in TMEM16A mRNA expression in the rice rat compared to the mouse (Figure 3G), whereas CFTR mRNA was significantly decreased (Figure 3H). Because carbachol-induced Cl^−^ secretion requires activation of the muscarinic 3 receptor (CHRM3) (Ousingsawat et al., 2009), we also assessed its transcript abundance. No differences in mRNA expression were found (Figure 3I). Thus, these data suggested that increased expression of TMEM16A, but not CHRM3, likely explained the augmented Cl^−^ secretion induced by carbachol.

### Cultured rice rat airway cultures secrete fluid in response to carbachol and display exaggerated fluid expansion in response to apical 3.5% NaCl

Based upon our ion transport studies, we hypothesized that carbachol would elicit exaggerated fluid secretion in the rice rat airway compared to the piglet or mouse. To test this hypothesis, we measured the apical surface fluid meniscus (Figure S1A-C) using previously published methods (Harvey et al., 2011). Baseline volumes were consistent with previously published values (Figure 4a-c) (Harvey et al., 2011). In the rice rat, basolateral carbachol significantly increased fluid secretion (Figure 4C). Carbachol did not elicit robust fluid secretion in the piglet (Figure 4B) or mouse (Figure 4A). Normalizing the fluid secretory response to baseline for each species illustrated a qualitatively greater response to carbachol in the rice rat relative to the mouse or pig (Figure 4D). However, a lack of a statistically significant interaction (F_6,39_ = 2.1, p = 0.08) precluded any post hoc analyses.

**Figure 4.**
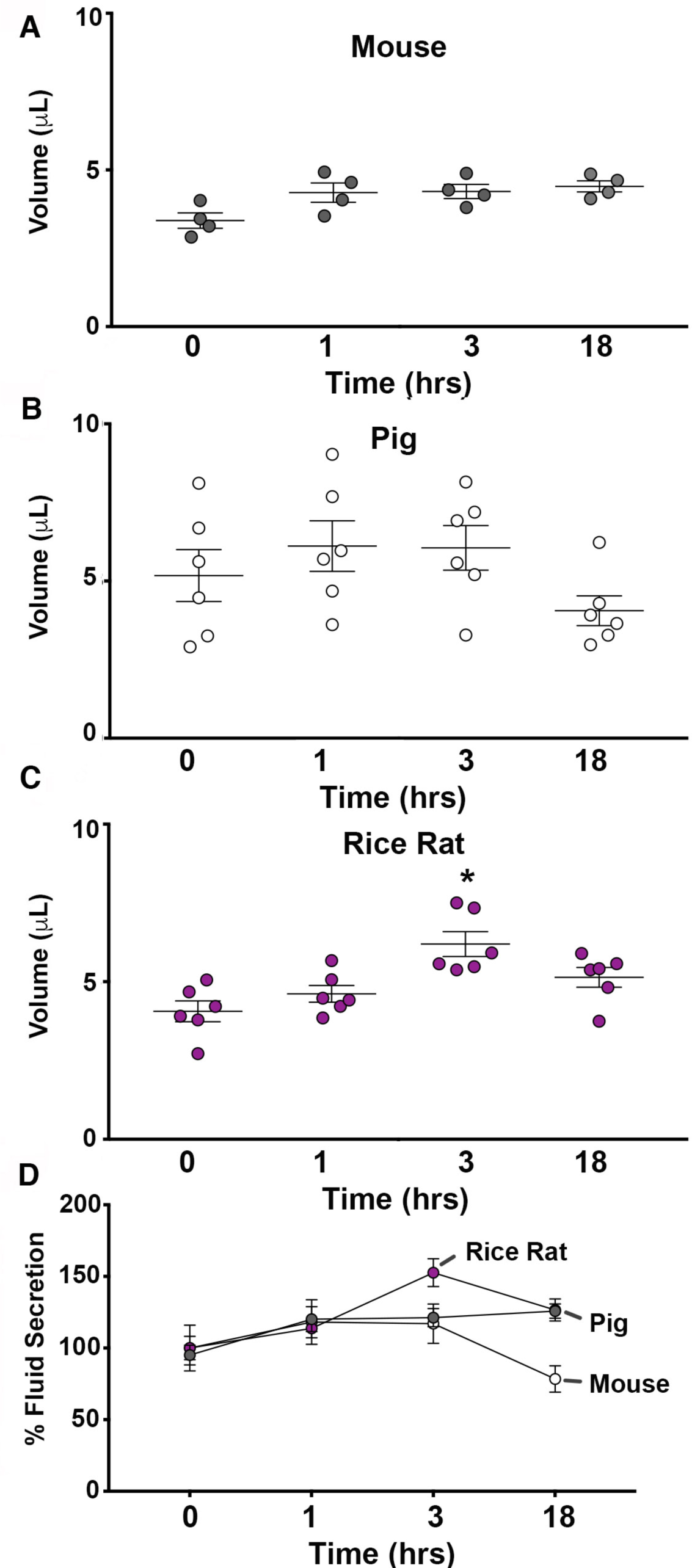
Rice rat primary cultures secrete fluid in response to carbachol. Apical fluid secretion in response to 100μM basolateral carbachol in mouse (A), pig (B), and rice rat (C) airway cultures. D) Fluid secretion responses normalized to baseline values and expressed as a percent for each species. Abbreviations: hour (hr). For all panels, n = 4 cultures from mice (2 female cultures and 2 male cultures representing 6 female and 6 male subjects); n = 6 cultures from the rice rat (3 female cultures, and 3 male cultures representing 6 female and 6 male subjects); n = 6 porcine cultures (3 female and 3 male representing 3 individual female and 3 individual male subjects). * = p < 0.05 compared to baseline. All data are shown as mean ± SEM.

We also challenged airway cultures with 3.5% hypertonic NaCl to mimic contact with a stimulus that might be encountered in the salt marsh (e.g., sea water aerosol) (Millero et al., 2008) (Figure 5A-D). In rice rat airway cultures, 3.5% NaCl increased the apical fluid at 2, 4, and 6 hours post challenge (Figure 5C). In contrast, apical fluid was only augmented at 1 hr in the pig cultures (Figure 5B), whereas the mouse showed insignificant augmentation (Figure 5A). Normalization of the fluid expansion relative to baseline for each species further highlighted greater expansion in the rice rat compared to the mouse or pig (Figure 5D).

**Figure 5.**
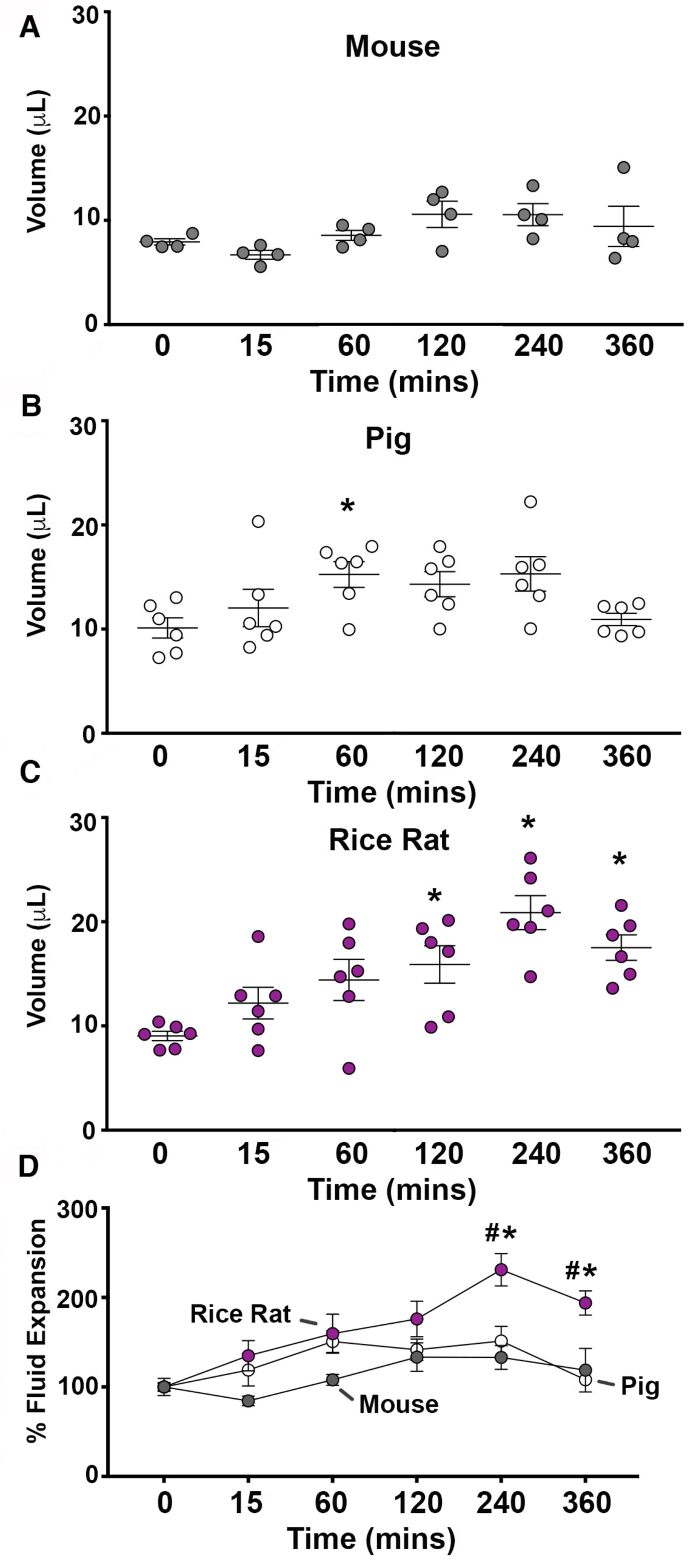
Rice rat primary cultures display and exaggerated fluid expansion to hypertonic saline. Surface fluid expansion in response to apical 3.5% NaCl increased in in mouse (A), pig (B), and rice rat (C) airway cultures. D) Fluid expansion responses normalized to baseline values and expressed as a percent for each species. Abbreviations: minutes (mins). For all panels, n = 4 cultures from mice (2 female cultures and 2 male cultures representing 6 female and 6 male subjects); n = 6 cultures from the rice rat (3 female cultures, and 3 male cultures representing 6 female and 6 male subjects); n = 6 porcine cultures (3 female and 3 male representing 3 individual female and 3 individual male subjects). For panels b-c* = p < 0.05 compared to baseline. For panel d, * = p < 0.05 compared to mouse; # p < 0.05 compared to pig. All data are shown as mean ± SEM.

## Discussion

We tested the hypothesis that the rice rat airway might have developed unique adaptations for the regulation of ASL. First, we investigated ion transport in rice rat, mouse and pig airway cultures. We found distinct ion transport properties, including minimal basal short circuit current, decreased amiloride-sensitive current, robust carbachol-mediated current, and minimal F&I-mediated current, in the rice rat airway. These findings suggested greater dependency upon CaCC, which was corroborated by finding increased mRNA expression of TMEM16A and decreased mRNA expression of CFTR. Moreover, we found that carbachol induced apical fluid secretion in rice rat airway cultures, whereas mouse and pig airway cultures were unaffected. Rice rat airway cultures also showed an exaggerated expansion of the ASL in response to 3.5% NaCl compared to the mouse and pig. Thus, these findings support the hypothesis that the rice rat airway exhibits unique ion transport mechanisms compared to animals that do not inhabit salt marshes.

We found minimal basal Isc in the rice rat airway cultures. Low basal Isc has been reported in the colon epithelium of mice with genetic disruption of α ENaC (Barker et al., 1998) and in intestinal segments from mice with targeted disruptions to the *CFTR* gene (Grubb, 1997). Consistent with that, we found limited amiloride-sensitive current and attenuated expression of ENaC in the rice rat airway culture compared to the mouse. This data and others suggest the rice rat is unique compared to other rodents. For example, in freshly excised Sprague Dawley rat tracheas, the amiloride-sensitive current is approximately ∼50-60 μA/cm^2^ (Tuggle et al., 2014), whereas in freshly excised mouse tracheas, it has been reported to be ∼15-20 μA/cm^2^ (Grubb et al., 2001). Epithelial cultures from the Wistar rat show an amiloride sensitive current that is approximately 2.0 μA/cm^2^ (Hahn et al., 2017), which was nearly 3 times greater than the average amiloride-sensitive current that we observed in the rice rat cultures (0.59 μA/cm^2^).

Although rice rat tracheal cultures may require steroids for *in vitro* expression of ENaC (Champigny et al., 1994), our findings did not support this hypothesis, as we observed no effect of treatment with aldosterone and dexamethasone on ENaC mRNA expression. Thus, one explanation for the low amiloride-sensitive current in the rice rat is naturally occurring mutations that impact ENaC trafficking and/or function. Indeed, α ENaCa, a naturally occurring truncated isoform of α ENaC, fails to generate amiloride sensitive current when expressed in *Xenopus* oocytes (Li et al., 1995). We are cloning rice rat ENaC subunits to examine this possibility. If true, then it is conceivable that Na^+^ is absorbed through an ENaC-independent mechanism(s) (O’Brodovich et al., 2008).

The secretion of Cl^−^ in response to carbachol was greater in the rice rat compared to the pig and mouse. Based upon published studies, the degree of carbachol-mediated Cl^−^ secretion in the rice rat is also greater than expected for the rat. Specifically, in tracheal cultures from Wistar rats, the amount of carbachol-mediated Cl^−^ secretion is 8.5 μA/cm^2^ (Hahn et al., 2017), which is approximately half of what we observed in the rice rat (19.7 μA/cm^2^). Consistent with a greater response to carbachol, rice rat cultures also showed elevated expression of TMEM16A mRNA compared to mouse. Thus, it is likely that enhanced TMEM16A expression contributes to the enhanced carbachol-mediated Cl^−^ secretion in rice rat. It is also possible that other channels contribute (Rock et al., 2009).

We found Cl^−^ secretion through CFTR was significantly decreased in the rice rat compared to the pig, and qualitatively less than the mouse. Paralleling this finding, mRNA expression for CFTR was also decreased in rice rat cultures compared to mouse cultures. This diminished CFTR activity appears to have minimal negative effects in the marsh rice rat airway, as they show no overt signs of lung disease, even as carriers of hantavirus (Holsomback et al., 2013). In CFTR knockout mice, and in humans and rats with CFTR mutations, TMEM16A is thought to compensate for loss of CFTR (Billet and Hanrahan, 2013; Clarke and Boucher, 1992; Tuggle et al., 2014). Ergo, it is likely that TMEM16A also compensates in rice rats.

Our study has limitations and advantages. The *in vitro* nature of our study can be considered both a limitation and advantage. Although we were not able to study the *ex vivo* or *in vivo* properties of the airway, the *in vitro* culture system we used shows morphologic features (Karp et al., 2002), transcriptional profiles (Pezzulo et al., 2011) and ion transport properties similar to that observed *in vivo* (Itani et al., 2011). Moreover, the *in vitro* nature of our study allows for analysis of the specific contribution of the epithelia, without the influence of other components, such as submucosal glands, smooth muscle, cartilage, nerves innervating the airway, and vasculature.

It is also worth noting that we did not investigate other epithelia in other tissues. The rice rat is an emerging research model used to study spontaneous periodontitis and medication-related osteonecrosis of the jaw (Aguirre et al., 2012; Messer et al., 2019; Messer et al., 2018). The interaction of the gingival epithelia with oral microbiota influences the status of health or disease (Tribble and Lamont, 2010). Although we did not investigate non-airway epithelia, it is interesting to consider that ion transport properties of other epithelia, such as the gingival epithelia, might also be distinct and/or modified compared to other animals. If so, then perhaps those properties contribute to the well-described predilection of periodontitis in the rice rat (Aguirre et al., 2017; Gotcher and Jee, 1981a, b; Gupta and Shaw, 1956a, b).

Previous studies have shown that epithelia from saline-adapted animals have decreased Na^+^ absorption and increased Cl^−^ secretion (Foskett et al., 1981; Kultz, 1993). If we interpret our data to indicate that the rice rat airway has evolved to also absorb less Na^+^ through ENaC and secrete more of Cl^−^ through CaCC, of what advantage might this be? We speculate that the rice rat airway when exposed to extreme heat (Silliman, 2014) and aerosols containing high NaCl concentrations (Gabriel, 2002), experiences a transient increased ASL tonicity. This increase in ASL tonicity activates the vagus nerve to elicit cholinergically-mediated Cl^−^ secretion (Wine, 2007), which acutely increases water on the airway surface. Simultaneously, the decreased Na^+^ absorption through ENaC and limited Cl^−^ absorption through CFTR ensure that the ASL expansion in response to increased tonicity is sustained. Combined, these processes likely help protect the airway against dehydration and ensure proper mucociliary transport (Matalon et al., 2015). However, having too much water on the surface of the airway is also of negative consequence as newborn mice lacking α ENaC die due to an inability to clear fluid out of the lungs (Hummler et al., 1996). Interestingly, we recently reported that atropine, a cholinergic antagonist, was required for proper respiratory patency during specific anesthesia regimens in the rice rat (Jiron et al., 2019). Thus, the unique airway transport of the rice rat might be advantageous in the salt marsh, but potentially disadvantageous in other environments.

In summary, our findings suggest that the rice rat airway has distinct ion transport properties that likely facilitate survival in the marsh salt environment. Perhaps examining the airway physiology of additional mammals that inhabit extreme environments might reveal similar, or perhaps even dissimilar, discoveries.

**Table 1.**
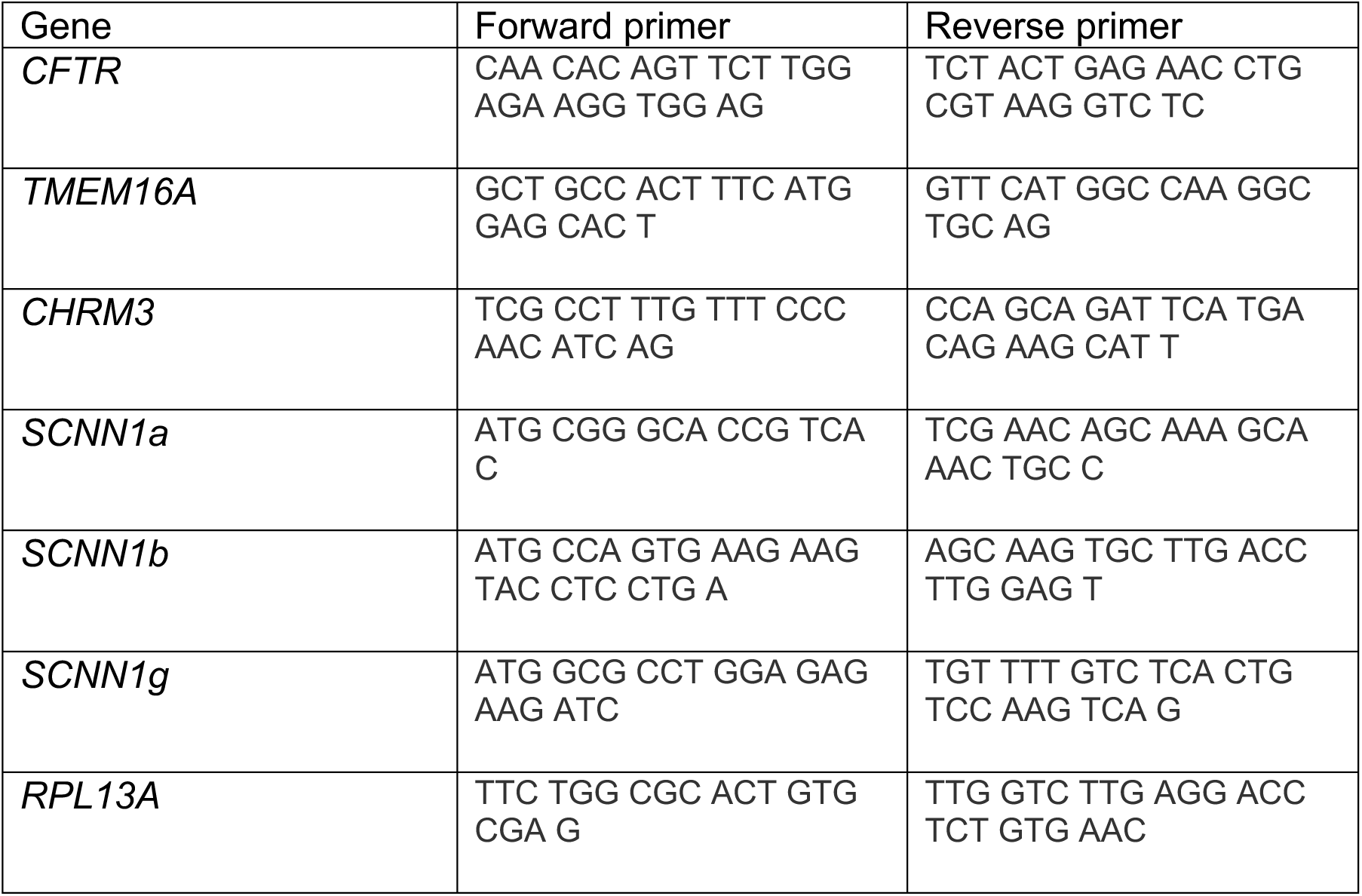
Primers used for cloning small coding regions of rice rat genes.

**Table 2.**
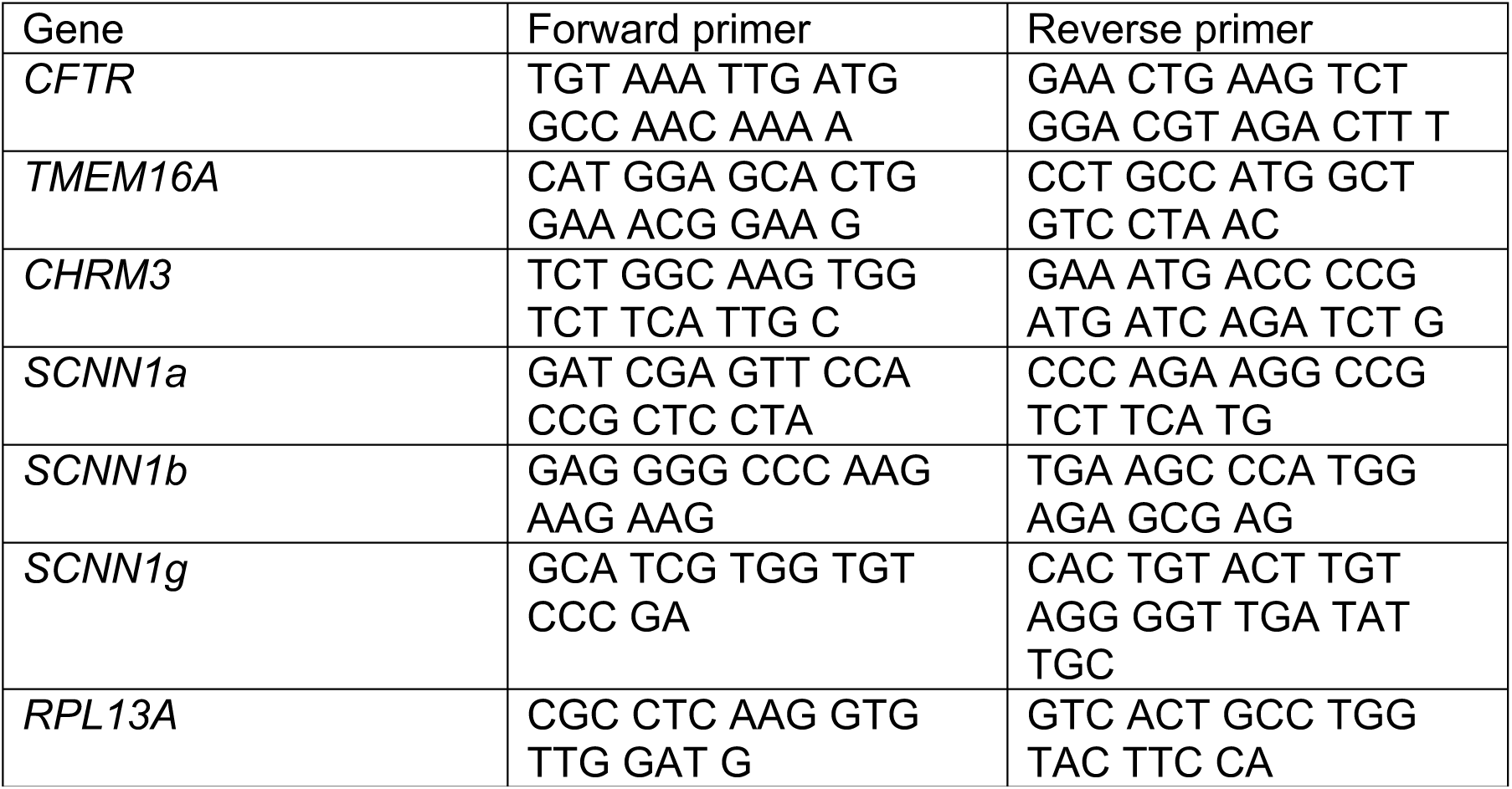
Primers used for quantitative RT-PCR.

## Acknowledgments

We thank Joshua Dadural, Emily Collins, Maria Guevara, Evelyn Castillo, Kalina Atanasova, Abel Abraham, Niousha Ahmar, Jacqueline Watkins, and Jessica Jiron for excellent technical assistance. We thank Dr. Jianyang Du and Dr. Mahmoud Abou Alaiwa for helpful suggestions and advice. GlyH-101 was a generous gift from the Cystic Fibrosis Foundation Therapeutics and R. Bridges.

## Author Contributions

LRR, YSJL, KMD, SPK, JIA, JZ, and JGM participated in the conception and design of the research. LRR, YSJL, KMD, SPK, JIA, JZ, and JGM performed the experiments. LRR, YSJL, KMD, SPK analyzed the data. LRR, YSJL, KMD, SPK, JIA, JZ, and JGM interpreted the results of the experiments. LRR and SPK prepared the figures LRR and SPK drafted the manuscript. All authors edited and reviewed the manuscript.

## Ethics statement

All studies were approved the University of Florida Institutional Animal Care and Use Committee.

## Materials and Methods

### Animal

Young male and female Landrace piglets (1 week old) (n = 5 female, n = 4 males) were purchased from an outside vendor and euthanized with intravenous euthasol (Henry Schein, Virbac) (Reznikov et al., 2013). Adult male (n =18) and female (n= 22) rice rats were derived from an in-house breeding colony (Aguirre et al., 2015) and euthanized by CO_2_ inhalation followed by cervical dislocation. Similarly, adult male (n = 8) and female C57BL/6 mice (n = 8) were obtained from an in-house breeding colony and euthanized by CO_2_ inhalation followed by cervical dislocation and bilateral thoracotomy. All procedures were approved by the University of Florida IACUC and the University of Florida Animal Care Services, which is an AAALAC-accredited animal care and use program.

### Primary cultures of differentiated airway epithelia

Epithelial cells were isolated from piglet tracheas by enzymatic digestion (Pronase, Roche; DNASE, Sigma) seeded onto collagen (Corning Collagen I, Rat Tail)-coated permeable filter supports (Corning Transwell polycarbonate membrane inserts, area = 0.33cm^2^, pore size = 4μM), and grown at the air-liquid interface using previously described procedures (Karp et al., 2002). Differentiated epithelia were studied at a minimum of 14 days after seeding. For the rice rat and mouse, 2-3 tracheas were pooled to isolate an adequate amount of cells for 3-4 permeable filter supports. For the pig, a single trachea yielded on average 24 inserts. Isolation procedures for rice rat and mouse airway epithelial cells were similar to those established for the piglet, except digestion of the tracheas occurred for only ∼24-30 hours for rice rat and mouse tracheas (compared to 48 hours for the piglet).

### Electrophysiological measurements of cultured airway epithelia

Cultured epithelia were studied in modified Ussing chambers (EasyMount Ussing Chamber System; Physiologic Instruments). Transepithelial voltage was maintained at 0 mV and short-circuit current (Isc) measured (VCC MC-8; Physiologic Instruments). Isc and delta (Δ) Isc were reported.

Cultured airway epithelia were bathed on the apical and basolateral surfaces with the following solution: 135 mM NaCl, 2.4 mM K_2_HPO_4_, 0.6 mM KH_2_PO_4_, 1.2 mM CaCl_2_, 1.2 mM MgCl_2_, 10 mM dextrose, 5 mM HEPES, pH 7.4 (NaOH). The solution was maintained at 37°C and gassed with compressed air (Chen et al., 2010; Stoltz et al., 2013).

The following protocol was performed: (a) measurements at baseline; (b) apical 100 μM amiloride to inhibit ENaC; (c) 100 μM basolateral carbachol to simulate CaCC (d) 30 μM CaCCinh-A01 to inhibit CaCC (He et al., 2011); (e) apical 10 μM forskolin and 100 μM IBMX to increase cAMP and activate CFTR; (f) apical 100 μM GlyH-101 to inhibit CFTR. All drug concentrations were selected according to our previous studies (Chen et al., 2010; Stoltz et al., 2013).

### Drugs used for Ussing chamber studies and cell culture

Ussing chamber studies: amiloride hydrochloride hydrate (Sigma); carbamylcholine chloride (carbachol) (Sigma); forskolin (Sigma); 3-isobutyl-1-methylxanthine (Sigma); Glyh-101 (a gift from the Cystic Fibrosis Foundation Therapeutics and Robert Bridges, Rosalind Franklin University of Medicine and Science, North Chicago, Illinois, USA); and CaCCinh-A01 (Sigma). Cell culture antibiotics (concentrations used are taken from reference (Karp et al., 2002): amphotericin B (Sigma); gentamicin (Gibco); penicillin-streptomycin (Gibco). Additional antibiotics/antifungals used include: tobramycin (Alfa Aesar), final concentration 25 μg/ml; fluconazole (Selleckchem), final concentration 2 μg/ml; piperacillin sodium (Selleckchem), final concentration 17.8 μg/ml; tazobactam (Selleckchem), final concentration 2.2 μg/ml. For examining a potential requirement of steroid hormones for expression ENaC, aldosterone (ACROS) was dissolved in 100% DMSO for a stock concentration of 1 mM. Stocks were diluted 1:1,000 in cell culture media for a final concentration of 1 μM. This dose was selected because it has been shown to increase ENaC expression in primary rat airway epithelial cells (Champigny et al., 1994). Dexamethasone (Fisher Scientific) was also dissolved in 100% DMSO to a stock concentration of 100 μM. Stocks were diluted 1:1,000 in cell culture media for a final concentration of 100 nM. Again, the dose was selected because it has been shown to increase ENaC expression in primary rat airway epithelial cells (Champigny et al., 1994). Cultures were incubated in drugs for 24 hours prior to harvest and examination of mRNA levels. This time point was selected based upon previous studies (Champigny et al., 1994; Sayegh et al., 1999).

### Immunofluorescence

Primary culture epithelial cells were fixed in 4% paraformaldehyde for 15 minutes. Following fixation, cells were washed three times for 10 minutes PBS and permeabilized for 15 minutes in 0.15% Triton X-100 (Fisher Scientific)/PBS solution. Cells were then blocked in a 4% normal goat serum (Jackson Immuno Research Labs)/Superblock buffer in PBS (ThermoFisher Scientific) for 60 minutes. Cells were incubated overnight at room temperature in primary antibodies. Following overnight incubation, cells were washed, incubated in appropriate secondary antibodies (listed below), and washed again. A Hoechst 33343 (ThermoFisher Scientific) stain was performed to identify nuclei. Cells were mounted in Vectashield Hardset (Vector labs) and cover slipped. Images were captured on a Zeiss Axio Zoom V16. Identical microscope settings for assessment of a single marker across species.

### Antibodies

Rabbit anti-ZO-1, 1:500 (Life Technologies, 617300); mouse anti-acetyl alpha tubulin, clone 6-11B-1, 1:500 (EMD Millipore, MABT868). The antigen for the ZO-1 antibody corresponds to residues 463-1109 of human ZO-1. This antibody has been used previously to identify ZO-1 in multiple species, including rat (Bordin et al., 2004) and human (Excoffon et al., 2009). According to the manufacturer, the antigen for the acetyl alpha tubulin is 15 S dynein fraction from the sea urchin sperm axoneme. This antibody has been used to identify human, *Drosophila*, and *Chlamydomonas* acetylated alpha tubulin (Piperno and Fuller, 1985). Secondary antibodies were used at 1:1000 and included: goat anti-mouse IgG (H+L) dylight 488 (ThermoFisher Scientific, 35505); alexa fluor 488 goat anti-rabbit IgG (H+L) (ThermoFisher Scientific, A11034).

### Fluid secretion studies

We implemented the reflected light microscopy method to measure the meniscus of the secreted fluid (Harvey et al., 2011). Briefly, cultures were placed on a heated stage maintained at 37*°*C and imaged using a Zeiss Axio Zoom V16. The following buffer bathed the basolateral side: 135 mM NaCl, 2.4 mM K_2_HPO_4_, 0.6 mM KH_2_PO_4_, 1.2 mM CaCl_2_, 1.2 mM MgCl_2_, 10 mM dextrose, 5 mM HEPES, pH 7.4 (NaOH). Light intensity and magnification were held constant for all cultures. A single image was taken at designated time points. Cultures were returned to the incubator in between imaging. The intensity profile of the image was examined using Zen Pro software analysis (Zeiss). The XY coordinates of the intensity profile correlated to distance (X) and intensity (Y).

The length of the fluid meniscus could therefore be readily identified as the distance between the greatest decrease in pixel intensity (at cell culture insert edge) minus the average baseline intensity (in the middle of the culture). A 4-point regressive curve used to convert meniscus lengths to volumes. The curve was generated by adding known volumes (5, 10, 15, 20 μl) of the bathing fluid were added to the apical surface of two separate cultures (e.g., duplicate standard curves) plotting regression curve (R^2^ value of 0.99: y = 1.5373e^0.001x). Apical fluid secretion to 100 μM basolateral carbachol was measured. The average salinity of sea water is 3.5% (Millero et al., 2008). Therefore, we also measured fluid secretion in response to 5 μl of apical 3.5% hypertonic saline (Harvey et al., 2011).

### RNA isolation, cloning and qRT-PCR

RNA from rice rat and mouse whole trachea, as well as airway epithelial cultures from rice rats and mice, were isolated using methods previously described (Reznikov et al., 2019). Briefly, a RNeasy Lipid Tissue kit (Qiagen) with optional DNase digestion (Qiagen) was used to isolate RNA. The RNA concentrations were assessed using a NanoDrop spectrophotometer (Thermo Fisher Scientific). RNA was reverse transcribed for trachea tissues (500 ng) and cultures (50 ng) using Superscript VILO Master Mix (Thermofisher) (Reznikov et al., 2019). RNA and master mix were incubated for 10 mins at 25°C, followed by 60 mins at 42°C, followed by 5 mins at 85°C.

Small segments of mRNA for rice rat CFTR, TMEM16A, α ENaC, β ENaC, γ ENaC, muscarinic 3 receptor (CHRM3) and RPL13a were amplified using primers designed to mouse orthologs from both cultured epithelia and tracheal epithelia (Table 1). Primers were ordered through IDT (Coralville, IA).

Amplification was achieved using Platinum Taq High Fidelity DNA polymerase (Thermofisher) and 50 ng of cDNA. Cycling parameters consisted of 35 cycles at 94 °C for 15 s, 52 °C for 20 s, and 68 °C for 30 s, and a final extension cycle of 3 minutes. Amplicons were run on a 1.5% agarose gel, extracted, and purified using a gel extraction kit (Qiagen). Amplicons were the sequenced using GENEWZ services (South Plainfield, NJ). Sequences were BLAST in NCBI and areas of 100% homology between rice rat and mouse were identified. Primers were designed according to areas exhibiting 100% homology to enable amplification of both mouse and rice rat genes with same primers (Table 2) and ordered through IDT (Coralville, IA). Transcripts for CFTR, TMEM16A, α ENaC, β ENaC, γ ENaC, and CHRM3 were quantified with qRT-PCR using RPL13a as a reference gene. All qRT-PCR data were acquired using fast SYBR green master mix (Applied Biosystems) and a LightCycler 96 (Roche). Standard ΔΔCT methods were used for analysis. SYBR green dissociation curves revealed the presence of single amplicons for each primer pair.

### Statistical Analysis

A one-way ANOVA was used to examine differences in ion transport. Post hoc comparisons between mouse and rice rat or pig and rice rat were achieved using Dunnett’s multiple comparisons test. A one-way ANOVA with repeated measures was performed to assess fluid secretion in mouse, rice rat or pig airway cultures. Post hoc comparisons were achieved using Dunnett’s multiple comparisons test for each species relative to baseline measures. For normalized secretion responses, a two-way ANOVA was performed with time as a repeated measure. When a significant interaction was observed, a Tukey’s multiple comparisons test was performed. Transcript abundances were assessed using a two-tailed unpaired student test (for each gene, mouse versus rice rat). Values with p < 0.05 were considered statistically significant. All analyses were performed in GraphPad Prism 7.0a.

**Figure S1.**
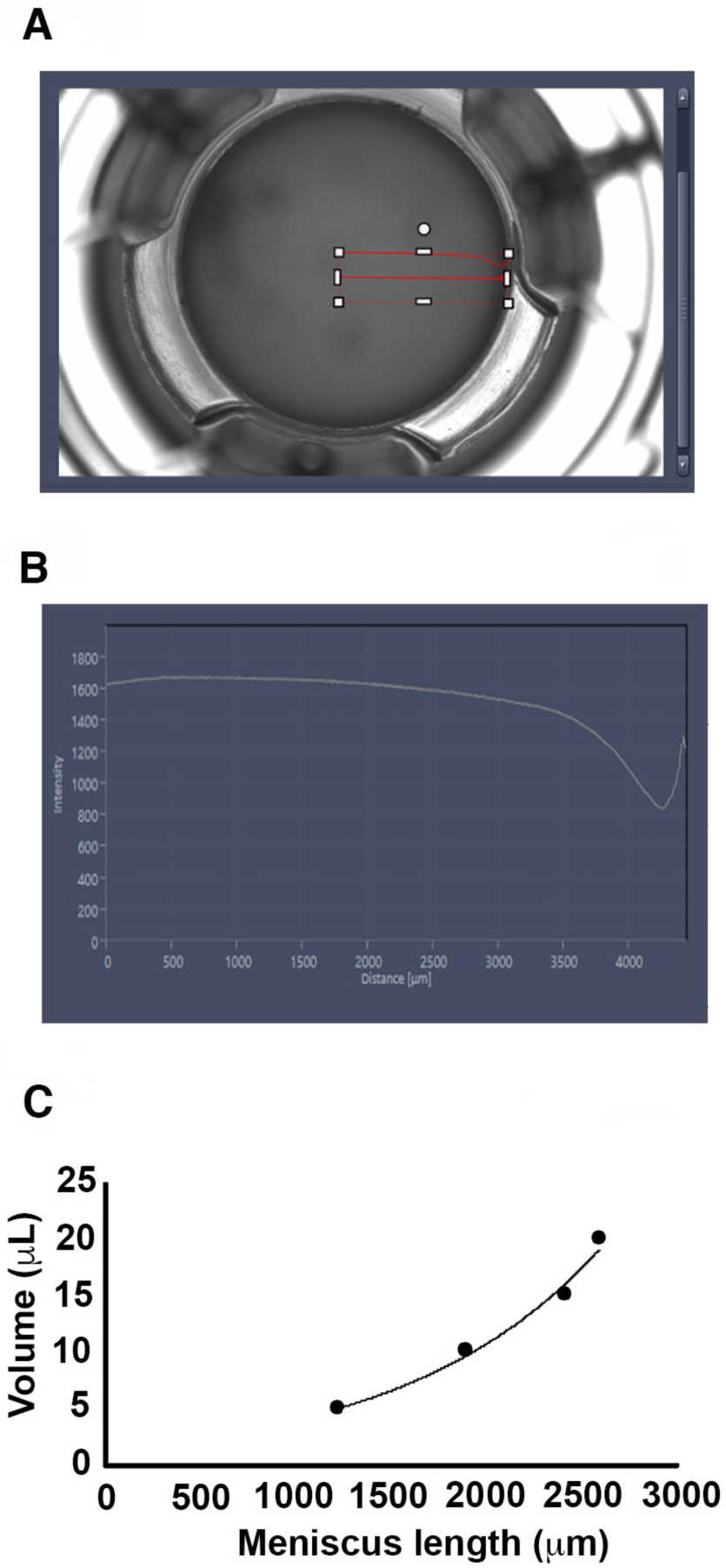
Measuring fluid secretion using reflected light and measurement of meniscus length. (A) Representative airway culture with intensity profile highlighted corresponding to red arrow shown directly above as red line with a “valley”. (B) Closer view of the red trace in panel A showing the XY coordinates corresponding to profile intensity (Y) and length (X). The length of the fluid meniscus was calculated as the distance between the greatest decrease in pixel intensity (at cell culture insert edge, shown on the right) minus the average baseline intensity taken from the middle of the culture. (C) Standard curve generated from 2 separate cultures when known volumes of fluid were added to the apical surface and the meniscus length measured. An exponential curve was fitted with a R^2^ value of 0.99 resulting in the following equation: y = 1.5373e^0.001x. This equation was used to extrapolate volume based upon meniscus length.

